# Rendering protein structures inside cells at the atomic level with Unreal Engine

**DOI:** 10.1101/2023.12.08.570879

**Authors:** Muyuan Chen

**Affiliations:** Division of CryoEM and Bioimaging, SSRL, SLAC National Accelerator Laboratory, Stanford University, Menlo Park, CA 94025, USA

## Abstract

While the recent development of cryogenic electron tomography (CryoET) makes it possible to identify various macromolecules inside cells and determine their structure at near-atomic resolution, it remains challenging to visualize the complex cellular environment at the atomic level. One of the main hurdles in cell visualization is to render the millions of molecules in real time computationally. Here, using a video game engine, we demonstrate the capability of rendering massive biological macromolecules at the atomic level within their native environment. To facilitate the visualization, we also provide tools that help the interactive navigation inside the cells, as well as software that converts protein structures identified using CryoET to a scene that can be explored with the game engine.

**Graphical abstract:** 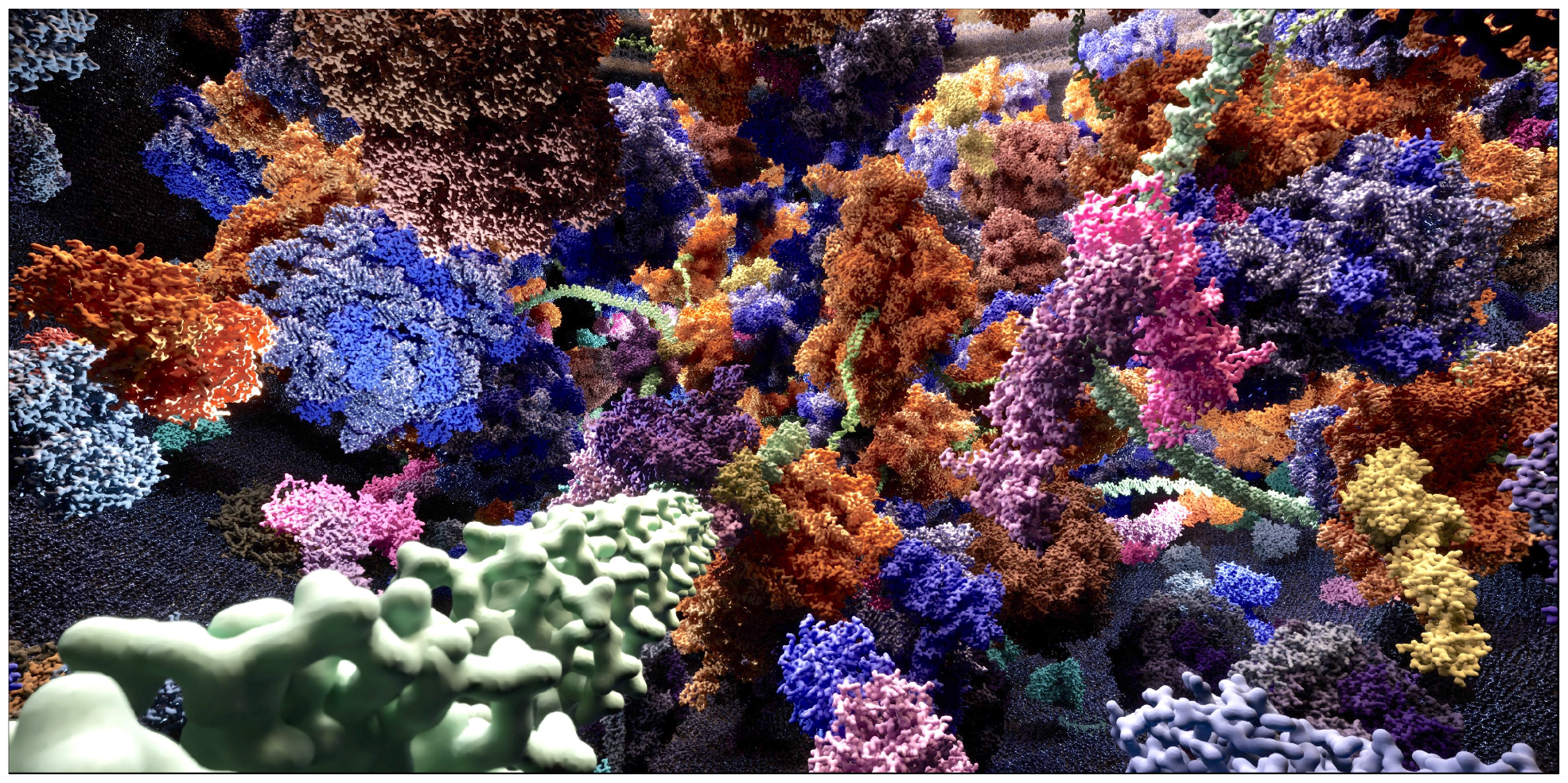

## Introduction

With the recent advances in cryogenic electron tomography (CryoET), structural studies of large macromolecules inside cells have become feasible. Combined with high-resolution structure available at protein databank (PDB) ^1^ and sequence-based structure prediction techniques, it is now possible to identify proteins of various types from the tomograms based on their structural features^2,3^. Location, orientation, and sometimes conformation of the target protein can be determined at their native state directly from the CryoET data^4,5^, providing in-depth views of the macromolecular world inside cells.

As more data is generated by the advanced experimental methods, the complexity of the cell revealed by CryoET also calls for new visualization techniques that can simultaneously display thousands of macromolecules inside the crowded cellular environment. Current visualization software, such as UCSF ChimeraX^6^, can only render one or a few protein structures at the atomic level. For cellular tomograms generated by CryoET, the data can only be rendered as slices of grayscale images or segmented volumes with low-resolution protein maps depicting their rough shapes^7^. The challenge of macromolecule rendering has become a bottleneck in many structural studies. For example, while near-atomic resolution ribosome structures of different conformational states have been obtained from tomograms of the bacterial and eukaryotic cells^8,9^, it is still highly challenging to directly visualize them at the determined resolution and interactively explore the cellular environment in three-dimension. The lack of visualization tools makes it harder for researchers to gain an intuitive and comprehensive view of the biological system. While some information, such as spatial correlation of molecules can be deduced statistically^4^, direct interactive visualization is still the most helpful in investigating the interaction between macromolecules inside cells and understanding the mechanisms of biological processes.

The challenge of macromolecule display inside cells is largely caused by the large number of triangles needed to render protein structures at high resolution. For example, while high-quality mesh of a human body typically requires less than 100,000 triangles, it takes nearly 300,000 triangles to render a single ribosome with a surface view at atomic resolution. Even a small scene of a few proteins can contain tens of millions of triangles, and drawing every individual one of them at every frame can easily exceed the capacity of modern GPUs. To overcome this issue, here we take advantage of a video game engine, Unreal Engine 5 (UE5), and demonstrate its capability of rendering large numbers of macromolecules at atomic resolution simultaneously and interactively.

Compared to the previous versions, UE5 features a new virtualized geometry system called Nanite. The Nanite system renders objects at different levels of detail automatically based on the distance of each object from the viewer, so the number of triangles needed to display at any given moment is drastically reduced. This makes it possible to render billions of triangles simultaneously with consumer-level GPUs. As a game engine, it also offers an interactive experience, so players can navigate inside the scene with a user-friendly interface in real-time. The latest versions of the game engine also feature the Python scripting capability, which provides a convenient way to connect it to external pipelines. With customized scripts, protein structures that are imported into the game can be automatically placed at their correct position determined from experimental data.

In this manuscript, we use two biological systems as examples to demonstrate the capability of UE5 rendering. To help users navigate the complex cellular scene and annotate macromolecular features, we develop in-game blueprints that enhance user control and interfaces. Finally, we also provide software tools that convert the location, orientation, and conformation of proteins from real CryoET datasets to a video game scene for interactive rendering, by mapping the high-resolution structures to their determined coordinates in the tomograms.

### Examples of biological systems rendering

To demonstrate the capability of the system, our first example is a salmonella minicell infected by P22 bacteriophage (**Figure 1**, Supplementary video 1). The scene is inspired by real CryoET tomograms from ^10^. From this dataset, many bacteriophages can be seen surrounding a small bacteria cell, with some P22 already anchored on the outer membrane of the cell, injecting their genome into the cytoplast. Numerous ribosomes, as well as densities of smaller proteins can be seen inside the cells. In this rendering, we estimate the size of the cell and the number of P22 and ribosomes from the CryoET data. From a minicell of ∼500 nm in diameter, we show the infection process of the P22, transcription and translation of the virus genome, as well as the respiration chain on the inner surface of the bacterial inner membrane.

**Figure 1.**
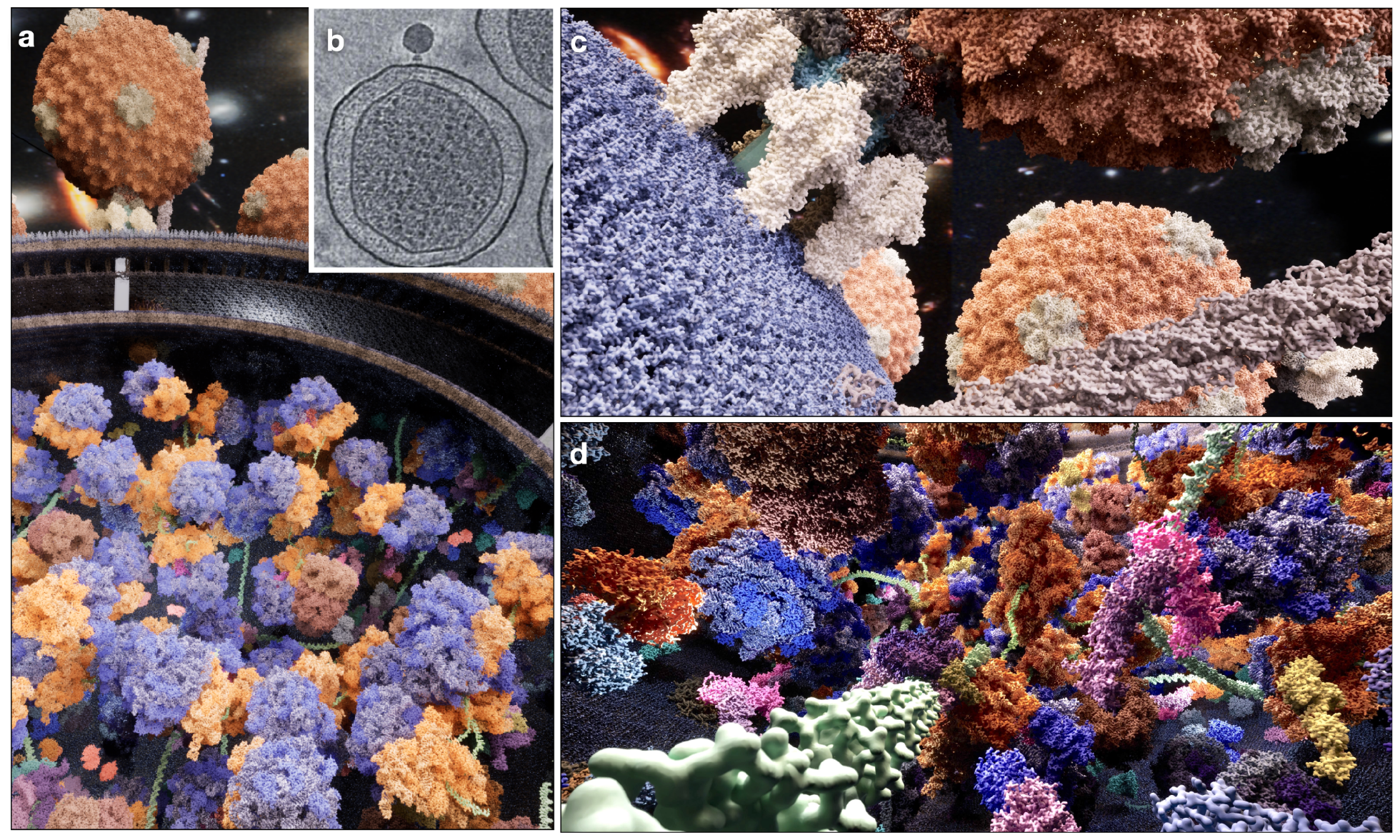
UE5 rendering of a salmonella minicell infected by P22 bacteriophages. (a) Overview of the scene. (b) Slice view of a reference tomogram from ^10^. (c) View outside the bacteria, showing viruses attached to the bacteria outer membrane and pilus extending out of it. (d) View inside the bacteria, showing virus DNA being transcribed and translated by the bacteria.

In this scene, we rendered 30 types of macromolecules with 580 different pieces of meshes, including the virus, bacterial flagella and pili, ribosomes, various chaperones, as well as proteins responsible for the glycolysis and respiration process. While the larger macromolecules, such as the viruses and ribosomes, can be determined from the CryoET data, many of the smaller proteins cannot be identified directly with the current technology. The list of those proteins were obtained from the proteomics data of Salmonella ^11^, and placed randomly inside the scene. The structure of each molecule is obtained from the PDB, and the mesh is generated using the surface view of the model, simulated at 3Å resolution. In sum, there are more than 40,000 objects, composed of around 1.5 billion triangles in the entire scene, and they can be rendered in real-time with a consumer GPU (a Nvidia RTX 4070 GPU is used in all examples).

In the second example, we explore the limit of the game engine’s power by rendering a piece of a eukaryotic cell, which includes parts of the Golgi `apparatus` and the chloroplast from a Chlamydomonas cell (**Figure 2**, Supplementary video 2). The scene is constructed using CryoET data from ^12,13^ as references. From CryoET data, the stacking of lipid membranes, as well as the protein coating the vesicles can be seen in the Golgi apparatus. Inside the chloroplast, thylakoid membranes stack on each other, and photosystem complexes are embedded in the membranes. In rendering this scene, we focus on the processes of photosynthesis and carbon fixation, which happen on the thylakoid membrane and inside the chloroplast lumen.

**Figure 2.**
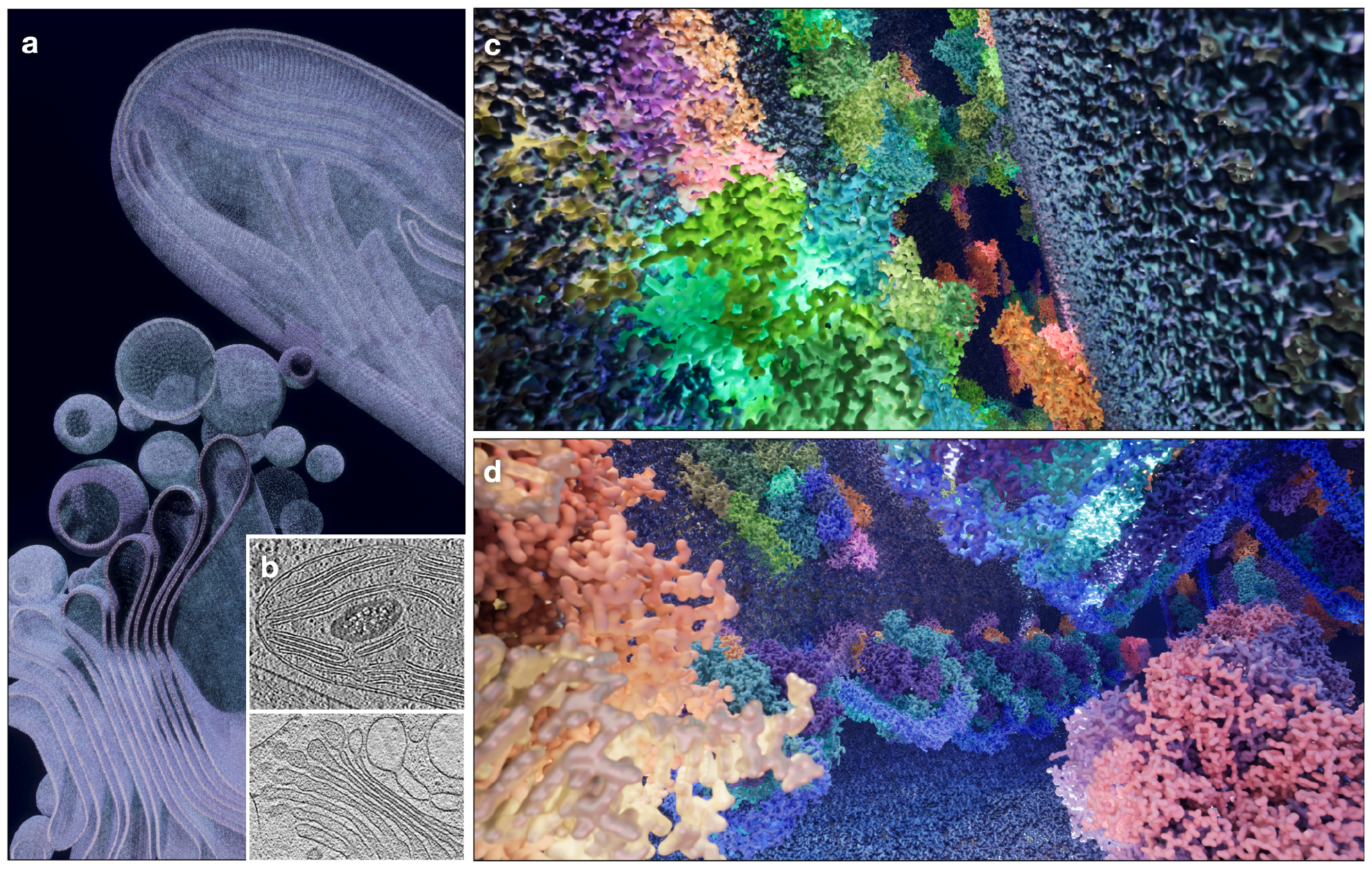
UE5 rendering of parts of the Golgi `apparatus` and the chloroplast from a Chlamydomonas cell. (a) Overview of the scene. (b) Slice view of reference tomograms from ^13,21^. (c) View inside the lumen of the thylakoid membranes, showing the photosystems and light harvesting complexes. (d) View of the chloroplast stroma, showing ATP synthase, RubisCO and ribosomes.

One of the main challenges in constructing the eukaryotic cell is the complexity of its membrane systems. Here, we programmably generate parametric descriptions of the membrane shapes and surface vectors and import the coordinates into the game engine. By piecing together many small square patches of lipid (from CHARMM archive^14^) at the corresponding location and orientation, we can build large organelles of various shapes. In the entire scene, which is roughly a 1μm cube in size, there are more than 20 million lipid molecules, each rendered at 3Å resolution. To render the lipid, as well as the protein complexes embedded on the membrane, we include more than 130,000 static mesh objects in the scene, which is made up of 11.6 billion triangles. Rendering of this scene is still well within the power of UE5 using a consumer GPU, and the player can navigate inside the scene to interact with the objects in real time.

### Tools for streamlined macromolecule placement

In classical game design, developers need to construct 3D models for each object, and manually place the models to build a scene. When the identity of macromolecules, as well as their location and orientation can be determined from a tomogram, the coordinates from CryoET data processing can be used to facilitate automatic object placement in the game engine.

Here, we developed a protocol that imports the high-resolution structures from the PDB into UE5, converts their position determined from the tomograms to the game engine coordinates, and places those molecules inside the cell using customized Python scripts.

To import the protein structures to UE5, we first fetch individual protein models from the PDB and display them in UCSF ChimeraX^6^. From ChimeraX, we can export the mesh of the models to GLB format, which can be read by the game engine. In the examples shown in the paper, we segment each protein by peptide chains and use surface rendering of each chain at 3Å resolution, but other forms of display, such as atoms-bonds or ribbons are also possible. After importing the GLB files to the game engine, Nanite needs to be enabled for each piece of mesh, and the color of the mesh can be adjusted by modifying its texture in UE5.

A few different approaches are available for placing the molecule meshes in the game scene. For macromolecules from real CryoET datasets whose orientations are determined through subtomogram averaging, we provide tools that convert the refinement results in EMAN2^15^ to the position of each particle in the game engine. To demonstrate the protocol, here we use the placement of spike proteins of NL63, a coronavirus, as an example^16^. From a tomogram (**Figure 3a**), we select the particles and determine the orientation of the crown of the spike, as well as the stalk that connects the spike to the membrane. Using existing tools provided in EMAN2, averaged structures can be mapped back to the original tomogram in their corresponding positions (**Figure 3b**). When the molecular model of each averaged structure is provided, we can now directly construct a scene in UE5, with every particle rendered at atomic resolution in their correct position and conformational state (**Figure 3c**, membrane not included). For protein complexes with symmetry, we also provide scripts that duplicate mesh objects and place them at the corresponding asymmetrical unit positions. Therefore, users only need to import the mesh objects of one asymmetrical unit to render the entire molecule.

**Figure 3.**
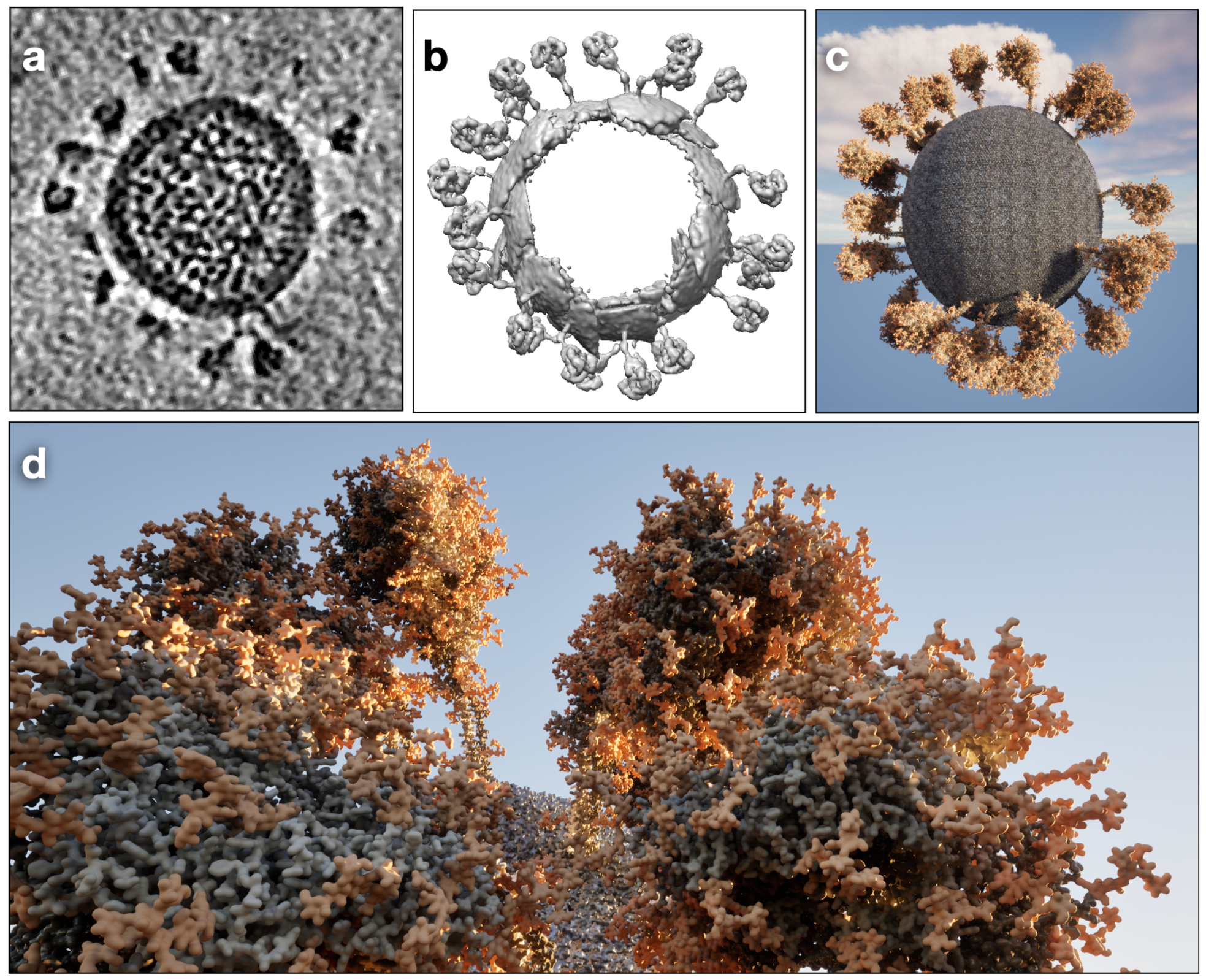
Converting real CryoET data to a UE5 scene. (a) Slice view of a tomogram of NL63 from ^16^. (b) Subtomogram averages of the spike proteins at different conformations mapped back to the tomogram, rendered in UCSF Chimera. (c) Atomic models of the same spike protein particles mapped to their corresponding positions in a UE5 scene. (d) Zoomed-in view of the UE5 scene.

Additionally, special protocols were developed for the placement of protein filaments with helical repeating patterns. To manually place a long filament in the game scene following any arbitrary path, we first build a trajectory using the sequencer tool, which controls the movement of objects during the recording of a movie. The in-game coordinates along the trajectory are then exported to a text file, which can then be read by a Python script that places the filament segments at the correct location and rotation along that path. For filament traced from tomograms, we also provide tools that convert the results from the filament tracing tool in EMAN2, or any path saved in the format of C-alpha protein backbone in a PDB file, to the coordinate system of the game engine.

### Blueprints for interactive cell visualization

Once a scene is constructed, users can move inside the scene and visualize the microscopic world around them. To better navigate the cellular environment, we also provide in-game protocols, called blueprints in UE5, that help control the movement of the character, as well as the interaction between the character and the environment.

Instead of having a fixed viewport and rotating the entire scene like the default mode in many structure visualization software, in UE5 the user controls a character, which is placed inside the scene. The game engine supports movement control with mouse and keyboard, as well as various types of controllers. Our blueprints add additional functionalities, including turning on/off a headlight, increasing movement speed, and toggling pass-through-wall mode, to make it easier to move around in a crowded cell.

One critical functionality for interactive visualization inside cells is to annotate each macromolecular object during the gameplay. To facilitate this, we introduce an additional piece of user interface and use a database to keep a record of every imported protein model in the scene. A data table can be provided in the form of a JSON dictionary that includes the description of each model. Using the line trace function provided by the game engine, we constantly look for the first visible object at the center of the view during the interactive session, find its description in the database, and display the information on the side of the screen (**Figure 4**).

**Figure 4.**
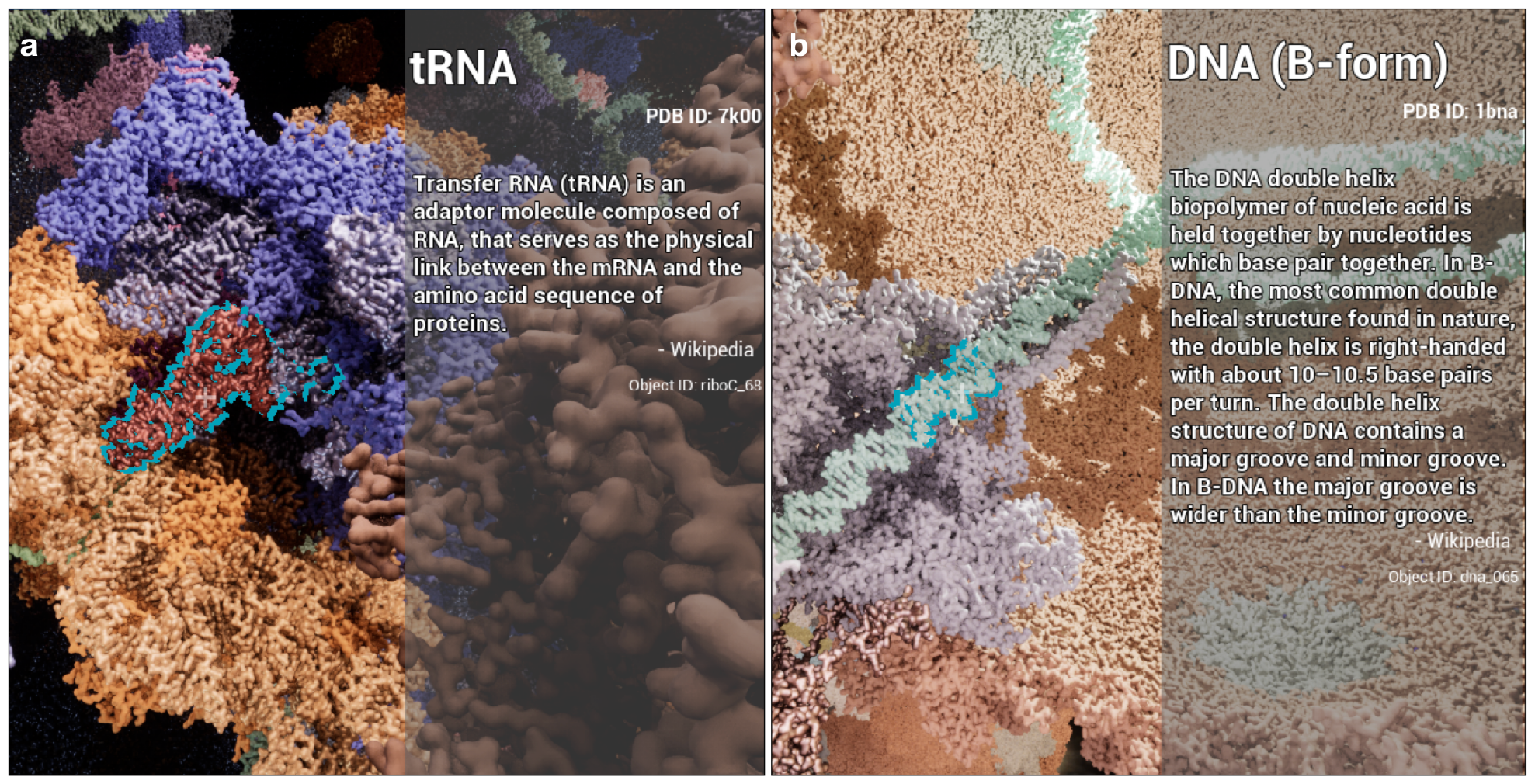
Annotation of macromolecular objects in the interactive game mode using UE5. (a) Annotation of a tRNA (highlighted in cyan outline) inside a ribosome. (b) Annotation of the DNA inside the portal of P22 virus, after some portal and capsid proteins were removed in the interactive mode.

Using a similar strategy, we also allow the player to “shoot” and destroy the first piece of model at the center of the view. Since each chain of a molecule is rendered as a separate object, this function provides a convenient way to “peel open” a complex and reveal the functioning site. For example, the player can remove the capsid and portal proteins of the P22 bacteriophage and directly visualize the trace of DNA inside the ejection machine of its tail (**Figure 4b**).

### Display of protein movement

To gain a thorough understanding of the macromolecular systems inside cells, it is critical to render the motion of proteins in addition to their static structures. Currently in UE5.3, the Nanite system only supports static meshes, i.e. the individual pieces of mesh cannot deform when Nanite is enabled. However, it is still possible to display the movement of proteins as the rigid body motion of the different parts of the molecule. When recording a video, the rigid body motion of the protein pieces can be modeled as the change of object transforms during the recording through the sequencer tool, similar to the control of camera movement. Alternatively, the conformational changes can also be rendered more intelligently by creating blueprint actors that include the mesh objects. For example, in the scene of the NL63 spike proteins shown in **Figure 3**, we created a blueprint for the spike proteins, and programed it so each spike can tilt continuously around a random axis. Additionally, when the tilt angle reaches the maximum allowed angle, or when the spike is colliding with another object, the tilt direction can be inverted to avoid unrealistic bending or clashing of molecules (supplementary video 3). The collision detection is also handled by the UE5, which automatically build convex hulls for the objects and send a signal each time two objects overlap.

Despite the power of the game engine, one major limitation in rendering protein movement is still our own lack of knowledge about the conformational dynamics of proteins at the atomic resolution, as well as the time scale it is happening. For example, while the structures of ribosomes with and without tRNA binding are known, it is unclear how the tRNA is inserted into its site without colliding with the rest parts of the structure. Therefore, in the two large scenes of the Salmonella and Chlamydomonas shown previously, we only present static structures in a “frozen” cellular environment. To display the movement of molecules in a realistic way, we are working on combining the UE5-based rendering system with the heterogeneity analysis methods for CryoEM/CryoET^17,18^, so that protein movement and conformational changes can be rendered at atomic level in the future.

### Support of virtual reality display

As the virtual reality (VR) technology develops, the interest of rendering the macromolecular world in VR devices is also increasing. While UE5 has good VR support in general, direct rendering of the massive scene of macromolecules in VR is still challenging. Even though the Nanite system has greatly reduced the resource requirement of the display, the resource required to render any of the scenes discussed above is well beyond the power of modern standalone VR devices. One possibility is to directly connect the VR device to a powerful enough computer that handles the rendering. However, a cable connection sacrifices the flexibility of many VR devices, greatly limiting the potential use cases. Additionally, a specialized control mode is needed to navigate the full 3D environment of the cell, which is still under development.

As an alternative approach, instead of visualizing the system interactively, we can also pre-generate stereoscopic (3D) panoramic (360°) videos and view them in VR devices. One simple way of making the 3D 360° video is to render the same 360° video twice, each time with a small, constant offset of camera locations (**Figure 5a** top). However, this approach preserves only the 3D view of the front part. If the user turns 180° wearing the VR device, the location of left and right eyes would be swapped, and the combined 3D view distorted. To overcome this, the cameras need to translate and rotate for each frame of the video (**Figure 5a** bottom). Since the recording of 3D 360° videos is not yet fully supported in UE5.3, we modified the built-in Movie Render Queue Additional Render Passes plugin to generate the videos. For each eye at each frame, the full view, which is 6k by 3k pixels in size, is divided into 60 tiles on a 10 by 6 grid, each rendered separately and stitched together afterwards (**Figure 5b**). Note that while the tiles merge almost seamlessly near the equator, some vertical stitching artifacts are still visible near the top and bottom of the view. Since the position of the two eyes are flexible when looking up or down, it is impossible to produce a true 3D video at those regions. The videos can be played through YouTube, or other software that display 3D 360° videos on VR devices.

**Figure 5.**
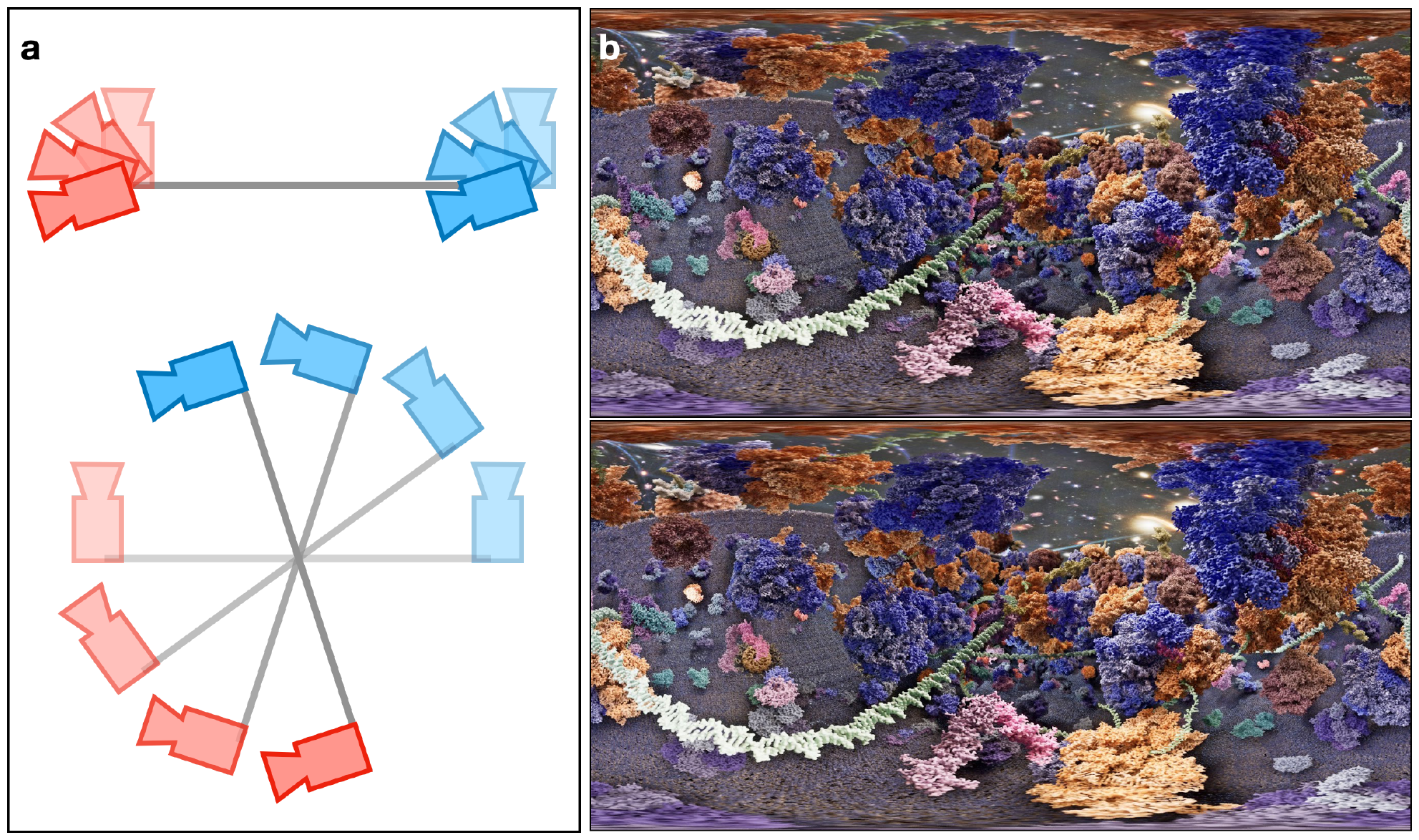
Rendering of 3D 360° videos. (a) Camera setup for each frame: top – fixed camera positions, bottom – rotating camera positions. (b) The panromantic view of the left (top) and right (bottom) eye view of one frame of the scene in Figure 1. A short 3D 360° video of the scene can currently be found at youtu.be/nqrMXAaOKYw

### Discussion and future work

In this work, we demonstrate the capability of rendering complex biological scenes with millions of molecules using a video game engine. The UE5 is free to download and free to use, until a certain amount of revenue is made using the engine (see EULA terms^19^ for details). All software tools developed in this work are freely available. Scripts for converting CryoET refinement results to game scenes are distributed through EMAN2, and a tutorial for constructing a UE5 scene using PDB structures is available at eman2.org/unreal_render.

While the power of game engine based rendering is already greatly exceeding the conventional methods for macromolecule display, there are still a number of improvements to be worked on in the future. First, although the placement of soluble proteins and filaments has been automated, the placement of membranes still requires customized programming and a parametric description of the shape of the organelles. In the future, we would also like to have automatic membrane construction from the segmentation of tomograms. In addition to identifying which tomogram voxels belong to a lipid membrane, this also requires the segmentation of disconnected pieces of membranes, the definition of the curvature and surface normal vector for each patch of membrane, as well as optimized smoothing to ensure the seamless connection between neighboring patches at the atomic level.

Second, much still need to be done to render the conformational changes of molecules. Since UE5 does not support Nanite for skeletal meshes yet, all conformational changes need to be handled as rigid body movement of individual pieces of static meshes. While we have shown the rendering of simple movement using UE5 blueprints, the process is far from straightforward. To address this, we are working on better integration between the CryoEM structural heterogeneity analysis protocol^17^ and the UE5 rendering scheme. To visualize realistic protein conformational changes in UE5, we aim to develop scripts that segment the protein into multiple meshes based on the pattern of movement, then automatically produce UE5 blueprints that allow the meshes to move along trajectories provided by the heterogeneity analysis of CryoEM data.

Additionally, we are also moving toward building more realistic protein-lipid interaction, so that membrane proteins can be correctly inserted into the lipid, instead of clashing with the lipid molecules. Currently, the membrane proteins are only placed inside the membrane, often occupying the same space as the lipid. To present the structure of membrane proteins more accurately, it would be ideal to take the information from the molecular dynamics (MD) simulation of membrane proteins and render each membrane protein with its surrounding lipid molecules at a reasonable conformation. As many membrane proteins rendered in the previous examples are large complexes, it may also be necessary to incorporate coarse-grain simulation methods^20^. With these features, we will be able to render the membrane systems in eukaryotic cells more realistically and visualize the protein-lipid interaction at the atomic level.

Finally, the interactive rendering of the biological systems at the atomic level can potentially be a great method for cell and molecular biology education. The high-quality videos produced by the game engine have been useful in presenting the biological systems, and the interactive game mode provides a new way to show students the marvel of the microscopic world.

Currently, we are still exploring various outreach possibilities, so the product of this work can be used as an educational tool that may inspire the next generation of cell and structural biologists.

## Acknowledgements

This work is supported by NIH grants R01GM150905 to M.C.

## Code availability

Scripts for converting particle locations in tomograms to mesh placement in UE5 are distributed through EMAN2, and a tutorial can be found at eman2.org/unreal_render. Some other scripts and notebooks for manipulating UE5 scenes are available at github.com/g5v991x/unreal5_protein_render. The interactive game, as well as the full UE5 projects that include the scenes discussed in the manuscript, are currently available upon request.

---

**Supplementary video 1**.

UE5 rendering of a salmonella minicell infected by P22 bacteriophages.

**Supplementary video 2**.

UE5 rendering of parts of the Golgi `apparatus` and the chloroplast from a Chlamydomonas cell.

**Supplementary video 3**.

Rendering of movement of NL63 spike proteins using UE5.

**Supplementary table 1.** Protein structures shown in the P22 infecting bacteria scene.

| Macromolecule name | PDB ID |
| --- | --- |
| tRNA | 7k00 |
| Ribosome | 7k00 |
| P22 tail spike | 1tyx |
| P22 portal | 1tyx |
| P22 ejection mechanism | N/A |
| P22 bacteriophage | 5uu5 |
| Hsp70/DnaK | 2kho |
| GroES | 1pcq |
| GroEL | 5w0s |
| ATP synthase | 6oqr |
| Cell wall | N/A |
| Bacterial lipoproteins | 1eq7 |
| Efflux pump | 5o66 |
| Bacteria flagella | 7cbl |
| NADH dehydrogenase | 4nwz |
| Outer membrane protein A | AF-P0A910 |
| Outer membrane protein C | 7jz3 |
| Type IVa pilus | 3jc8 |
| Respiratory complex I | 3m9s |
| Type III secretion system | 7ah9 |
| Ubiquinol oxidase | 1fft |
| DNA (B-form) | 1bna |
| Bacteria inner membrane | CHARMM |
| Bacteria outer membrane | CHARMM |
| DNA template | 7mkd |
| RNA polymerase | 7mkd |
| DNA template | 6ztj |
| Bacterial Expressome | 6ztj |
| Enolase | 1e9i |
| Phosphoglycerate kinase | 1zmr |
| Glycerol-3-phosphate dehydrogenase | 3da1 |
| HtpG (bacterial Hsp90) | 2ioq |
| Glyceraldehyde 3-phosphate dehydrogenase | 6utn |
| Glycerol kinase | 1bo5 |
| Malate dehydrogenase | 2pwz |

